# Overlapping Definitive Progenitor Waves Divide and Conquer to Build a Layered Hematopoietic System

**DOI:** 10.1101/2020.12.24.424302

**Authors:** Laina Freyer, Lorea Iturri, Anne Biton, Elisa Gomez Perdiguero

## Abstract

Adult innate immune cells are part of a layered hematopoietic system constructed from definitive hematopoietic stem and progenitor cells (HSPC) with diverse origins during development. One source of HSPC are fetal hematopoietic stem cells (HSC) that provide long-term reconstitution throughout life. However, the extent to which HSC produce mature cells *in utero* is only recently being uncovered. This is in part due to the added complexity of an overlapping wave of definitive progenitors that derive from yolk sac erythro-myeloid progenitors (EMP). HSC and EMP are generated from spatiotemporally distinct hemogenic endothelia, yet they both migrate to the fetal liver niche where they co-habitate and are presumed to reach their full potential. Delineation of the respective HSC and EMP pathways towards developmental immune cell differentiation has been confounded by challenges in ontogeny-specific cell labeling. In this study, *in vivo* inducible pulse chase labeling revealed that HSC contribute little to fetal myelopoiesis and that EMP are the predominant source of mature myeloid cells until birth. This is similar to what has been reported for the erythroid branch of hematopoiesis thereby establishing a developmentally-restricted privilege for erythro-myeloid differentiation from EMP compared to HSC. Tracing the origins of mature cells to the progenitor level by immunophenotyping and single cell RNA sequencing uncovered a dichotomy in the allocation of fetal liver EMP and HSC to myeloid progenitor subsets, both in timing and lineage bias. This has exposed an uncoupling between developmental granulopoiesis and monopoiesis from EMP and HSC pathways, and provides a framework for future studies of HSC-dependent and -independent hematopoiesis.

**HIGHLIGHTS:**

- EMP-to-HSC switch in fetal liver myelopoiesis occurs late in gestation
- EMP are efficient at producing early transit amplifying erythroid and myeloid intermediates
- scRNA-seq reveals three trajectories of EMP myelopoiesis
- Myeloid lineage commitment during development is cell type and ontogeny specific

## INTRODUCTION

Attempts to delineate mechanisms of developmental hematopoiesis are faced with many challenges brought on by the complexity of layering that has evolved to meet the needs of the growing embryo while simultaneously setting the groundwork for a lifetime of hematopoietic maintenance. The persistence versus resolution of different layers of the innate immune system (reviewed in Perdiguero & Geissmann, 2016) highlights the importance of exploring the link between developmental events and adult disease (Bennett et al., 2018; Honold and Nahrendorf, 2018; Loyher et al., 2018; Zhu et al., 2017). Furthermore, understanding how hematopoietic progenitor survival, expansion and differentiation is regulated with respect to ontogeny has further translational relevance to the production of immune cells *in vitro*.

The mammalian hematopoietic system is comprised of mature cells originating from at least two distinct waves of definitive progenitors, erythro-myeloid progenitors (EMP) and hematopoietic stem cells (HSC). EMP are generated from the blood island and vascular plexus of the extraembryonic yolk sac between embryonic days E8.25 and E11.5 (Frame, 2015; Kasaai et al., 2017; Lux et al., 2008; Palis et al., 1999). They are responsible for red blood cells that support embryonic survival until birth (Chen et al., 2011; Soares-da-Silva et al., 2021) and tissue resident macrophages and mast cells that persist into adult life and replenish independently of HSC (Ajami et al., 2011; Gentek et al., 2018; Ginhoux et al., 2010; Gomez Perdiguero et al., 2015; Hashimoto et al., 2013; Hoeffel et al., 2015; Schulz et al., 2012; Yona et al., 2013, 2013). Overlapping with EMP production is the wave of HSC that emerge intraembryonically (Ana Cumano et al., 1996; Cumano et al., 2001; Medvinsky and Dzierzak, 1996; Yvernogeau and Robin, 2017) from the aorta-gonad-mesonephros (AGM) region and other hemogenic sites (placenta, vitelline and umbilical arteries) between embryonic days E10 and E12 (De Bruijn et al., 2000; Gekas et al., 2005; Ottersbach and Dzierzak, 2005). HSC are defined by the ability for long-term multilineage reconstitution when transplanted to adults. Adding to the complexity of these overlapping waves are developmentally restricted HSC that specialize in tissue resident lymphoid output (Beaudin et al., 2016; Elsaid et al., 2020; Simic et al., 2020). Therefore there exist cells that are functionally important in adults but which are only produced during development.

The EMP origin of tissue resident macrophages and mast cells has been described (Gentek et al., 2018; Gomez Perdiguero et al., 2015; Hoeffel et al., 2015; Schulz et al., 2012), but the extent to which EMP and fetal HSC contribute to other mature hematopoietic cells during development and postnatal life has only recently begun to be revealed for erythroid and lymphoid lineages (Elsaid et al., 2021; Simic et al., 2020; Soares-da-Silva et al., 2021). Among myeloid cells, it is known that EMP are efficient at generating macrophages and megakaryocytes by direct differentiation within their niche of origin, the yolk sac (McGrath et al., 2015; Iturri et al., 2021). However, knowledge about the regulation of EMP expansion and differentiation once they have seeded the fetal liver is limited. EMP are the first to colonize the fetal liver, followed shortly thereafter by HSC (Kieusseian et al., 2012), and it is within that environment that they are thought to excel at producing committed intermediate progenitors responsible for transit amplification prior to differentiation.

The differentiation of HSC has long been characterized as a canonical hierarchical cascade of progenitor states based on expression of the cell surface markers CD16/32 and CD34. In this manner, a common myeloid progenitor (CMP) has been isolated from bone marrow that subsequently gives rise to further committed megakaryocyte-erythroid progenitors (MEP) and granulocyte-monocyte progenitors (GMP) (Akashi et al., 2000). While this hierarchy is phenotypically and functionally conserved in the E14.5 fetal liver (Traver et al., 2001), there were notable differences in burst activity in colony-forming unit (CFU) assays, and a distinction was not made between EMP- and HSC-derived progenitors due to the unavailability of lineage tracing tools at the time. Pulse chase labeling using *Tie2*^*MeriCreMer*^ suggest that HSC do not contribute significantly to GMP and MEP compartments until late in gestation, thereby leading to the hypothesis that they are predominantly of EMP origin (Gomez Perdiguero et al., 2015).

Identifying fetal immune cells that are of EMP or HSC origin is complicated by many factors. Inherent to EMP and HSC development are their endothelial-to-hematopoietic origins and co-habitation in the fetal liver. This has complicated efforts to distinguish progeny from one another due to many shared cell surface markers and therefore necessitates timed lineage tracing methodologies validated by multiple approaches. Pulse chase labeling has proven indispensable for tracing EMP independently of HSC using *Csf1r*^*MeriCreMer*^ mice (Gomez Perdiguero et al., 2015; Schulz et al., 2012). Recently, *Cdh5*^*CreERT2*^ mice have been used to label EMP or HSC by separating the timing of labeling to encompass endothelial cells that give rise to hemogenic endothelia at different stages (Gentek et al., 2018; Simic et al., 2020).

Due to the immunophenotypic similarities between EMP and fetal HSC, the mechanism(s) that distinguish them functionally remains an open question. Genetic manipulations and prolonged *in vitro* culture from pre-circulatory concepti support the notion that yolk sac EMP develop from functionally distinct hemogenic endothelia compared to intraembryonic sources of HSC, suggesting a role for cell intrinsic programming (Chen et al., 2011; Ganuza et al., 2018). Both EMP and HSC express the transcription factor MYB, although it is only required for survival of HSC and not EMP (Mukouyama et al., 1999; Schulz et al., 2012; Sumner et al., 2000; Iturri et al., 2021). Furthermore, HSC rely on blood flow hemodynamic forces and Notch signaling within the AGM whereas EMP are independent of these factors (Kasaai et al., 2017; Kumano et al., 2003; Sumner et al., 2000). While intrinsic factors are likely at play in functionally separating EMP from HSC, the impact of their secondary niche cannot be denied. The fetal liver provides a stromal microenvironment important for both structural support as well as a trophic secretion of growth factors utilized in HSPC expansion and commitment (Gao et al., 2018). EMP are highly sensitized to erythropoietin compared to HSC (Soares-da-Silva et al., 2021), indicating that a competition for limiting growth factors within the fetal liver niche may act as a driving force for differentiation. Whether the same holds true for myeloid progenitors remains to be determined.

In this study, we utilized *Cdh5*^*CreERT2*^ and *Csf1r*^*MeriCreMer*^ mice for *in vivo* pulse chase labeling of EMP and HSC. We found that HSC contributed minimally to myeloid cells during development, primarily monocytes followed by a late production of mast cells and neutrophils just before birth. Rather, the majority of fetal liver myeloid cells were of EMP origin until E16 and could still be found circulating in significant numbers postnatally. This prompted us to trace their origins at the progenitor level with respect to CMP, GMP and MEP classification. This revealed that the phenotypic CMP compartment in the fetal liver was almost entirely populated by HSC whereas EMP rapidly delegated to GMP and MEP intermediates. Around E15, an EMP-to-HSC switch in the production of GMP and MEP was observed followed one day later by a switch in the appearance of mature cells. The generation of mature myeloid cells from both EMP and HSC was lineage- and ontogeny-dependent with uncoupling of granulopoiesis and monopoiesis despite presumptions that they would be produced in an unbiased manner from the same GMP intermediates. This was supported by single cell RNA-sequencing (scRNA-seq) of EMP-derived progenitors in the fetal liver that were largely biased towards mast cell and granulocyte intermediates at E12.5. Underrepresentation of molecular signatures for Kit^+^ presumptive macrophage-producing myeloid progenitors that would be capable of locally giving rise to tissue macrophages suggested that commitment of these progenitors may occur mostly within the yolk sac. This is in line with observations that the intraembryonic trafficking of pre-macrophages peaks at E10.5 and is nearly complete by E12.5 (Stremmel et al., 2018). Therefore, the expansion of fetal liver tissue macrophages from E12.5 could be accounted for by local proliferation rather than input from fetal liver myeloid progenitors.

Collectively, we report that HSC are not a predominant source of fetal myeloid cells during development and that the *in vivo* execution of myeloid differentiation within the mouse fetal liver utilizes molecular programs that differ both in developmental time and lineage selection between EMP and fetal HSC.

## RESULTS

### HSC do not contribute significantly to mature myeloid cells until late gestation

To determine the level at which HSC contribute to mature hematopoietic cells during development, we analyzed *Cdh5*^*CreERT2*^ *Rosa26*^*YFP*^ embryos. A critical advantage of this system compared to the *Csf1r*^*MeriCreMer*^ *Rosa26*^*YFP*^ approach is that it bypasses lineage bias that may occur due to role of *Csf1r* in myeloid cells. By taking advantage of spatiotemporal differences in the generation of EMP and HSC from endothelia (Gentek et al., 2018; Simic et al., 2020), HSC were selectively labeled by injection of 4-hydroxytamoxifen (OHT) at E10.5, whereas EMP were labeled by injection of OHT at E7.5 with the caveat that this also labeled some precursors of HSC-generating endothelium (**Figure S1A-B**). To control for inherent variabilities between individual pulse chase labeling manipulations and to take into account the labeling of HSC by both injection timepoints, the percentage of YFP among mature cells was normalized either to the percentage of YFP among microglia (MG, CD45^+^ Kit^neg^ F4/80^hi^ CD11b^+^) for E7.5 OHT or to the percentage of YFP among LSK (Lin^neg^ Sca1^+^ Kit^+^) for E10.5 OHT (**Figure 1A-B**). Flow cytometry identified mature erythroid (red blood cell, RBC; megakaryocyte, Mk) and myeloid (mast cell, Mast; macrophage, MF; monocyte, MO; neutrophil, Nt) cells (**Figure S1C**).

**Figure 1.**
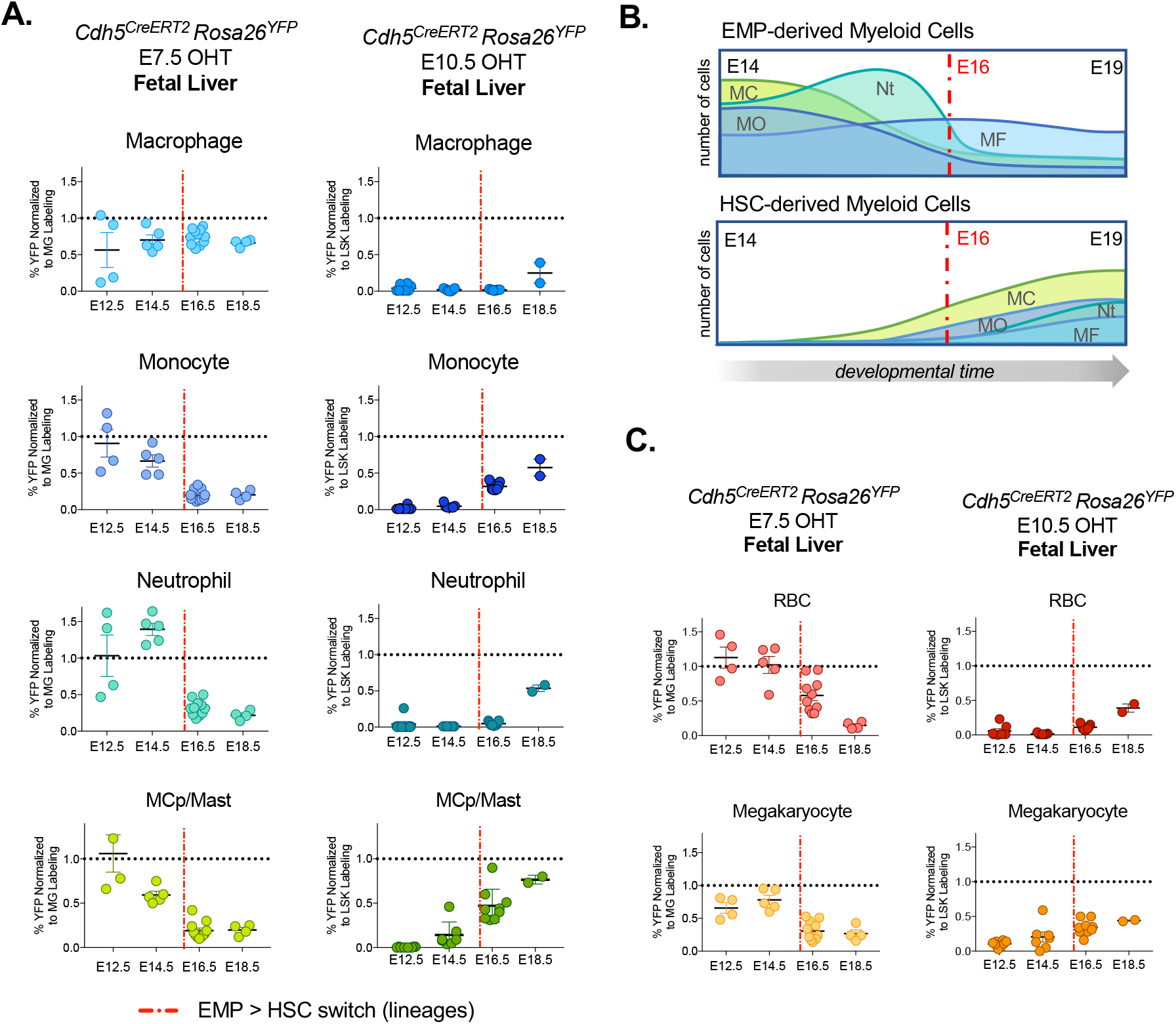
The ‘switch’ to HSC myelopoiesis occurs late and is lineage specific. **A.** Origins of mature myeloid cells during development as labeled by *Cdh5*^*CreERT2*^ *Rosa26*^*YFP*^ injection of 4-hydroxytamoxifen (OHT) at E7.5 (left, to label EMP) or E10.5 (right, to label HSC). For E7.5 OHT pulse labeling, the percent of YFP labeling was normalized to % YFP^+^ among microglia (CD45^+^ F4/80^+^, see Figure S1A). For E10.5 OHT, the percent of YFP labeling was normalized to % YFP^+^ among LSK (Lin^neg^ Sca1^+^ Kit^+^, see Figure S1A). Red dashed lines indicates the switch between EMP and HSC output at E16. **B.** Collective representation of EMP and HSC contribution to myeloid lineages from E14 to birth based on pulse chase labeling results. **C.** Origins of mature erythroid cells as labeled by *Cdh5*^*CreERT2*^ *Rosa26*^*YFP*^ injection of OHT at E7.5 or E10.5. YFP labeling was normalized using the same approach as in Panel A. MG, microglia; RBC, red blood cell; MCp, mast cell progenitor; MC, mast cell; MO, monocyte; Nt, neutrophil; MF, macrophage. Data are represented as mean ± SEM.

As expected, tissue macrophages (CD45^+^ Kit^neg^ F4/80^hi^ CD11b^+^) derived predominantly from EMP as reflected by the high percentage of YFP labeling that paralleled that of the microglia (**Figure 1A**, **Figure S1A**). The input from HSC to tissue macrophages, starting only at E18.5, was insignificant considering the total number F4/80^hi^ YFP^+^ macrophages per fetal liver. Furthermore, Kupffer cells retained postnatal labeling that was induced by E7.5 OHT injection *in utero* (**Figure S1B**). Other mature myeloid cells, such as monocytes (CD45^+^ Kit^neg^ F4/80^lo^ CD11b^+^ CSF1R^+^ Ly6C^+/−^), neutrophils (CD45^+^ Kit^neg^ CSF1R^neg^ CD11b^+^ Ly6G^+^) and mast cell progenitors (MCp) or mast cells (Lin^neg^ Sca1^neg^ Kit^+^ Itgb7^hi^ CD16/32^hi^) (Grootens et al., 2018) displayed an EMP-to-HSC switch around E16 (**Figure 1A-B**). Mature red blood cells (RBC, Lin^+^ CD45^neg^ Kit^neg^) and megakaryocytes (Lin^neg^ CD45^neg^ Kit^neg^ CD41^+^) also followed this pattern (**Figure 1C**). Notably, the normalization of labeling among macrophages and megakaryocytes following E7.5 OHT injection did not equilibrate with microglia labeling, supporting the observation that they begin differentiation locally in the yok sac niche at earlier developmental stages (McGrath et al., 2015; Iturri et al., 2021).

Taken altogether, EMP are responsible for generating the majority of tissue macrophages, mast cells and neutrophils during development whereas HSC myeloid output is largely limited to fetal monocytes and late waves of mast cells and neutrophils. The late (>E16) EMP-to-HSC switch in production of mature myeloid cells is also conserved for mature erythroid cells indicating that EMP are given privilege over HSC for erythro-myeloid differentiation during embryonic development.

### Late commitment of HSC to erythroid and myeloid progenitors

The lack of production of mature erythroid and myeloid cells from HSC in the fetal liver prompted us to examine the ontogeny of lineage commitment at the progenitor level. Using the paradigm of CMP to GMP/MEP commitment (Akashi et al., 2000; Traver et al., 2001), we examined cell surface of expression of CD16/32 and CD34 among lineage traced progenitors (LK, Lin^neg^ Kit^+^ Sca1^neg^) (**Figure 2A-B, Figure S2**).

**Figure 2.**
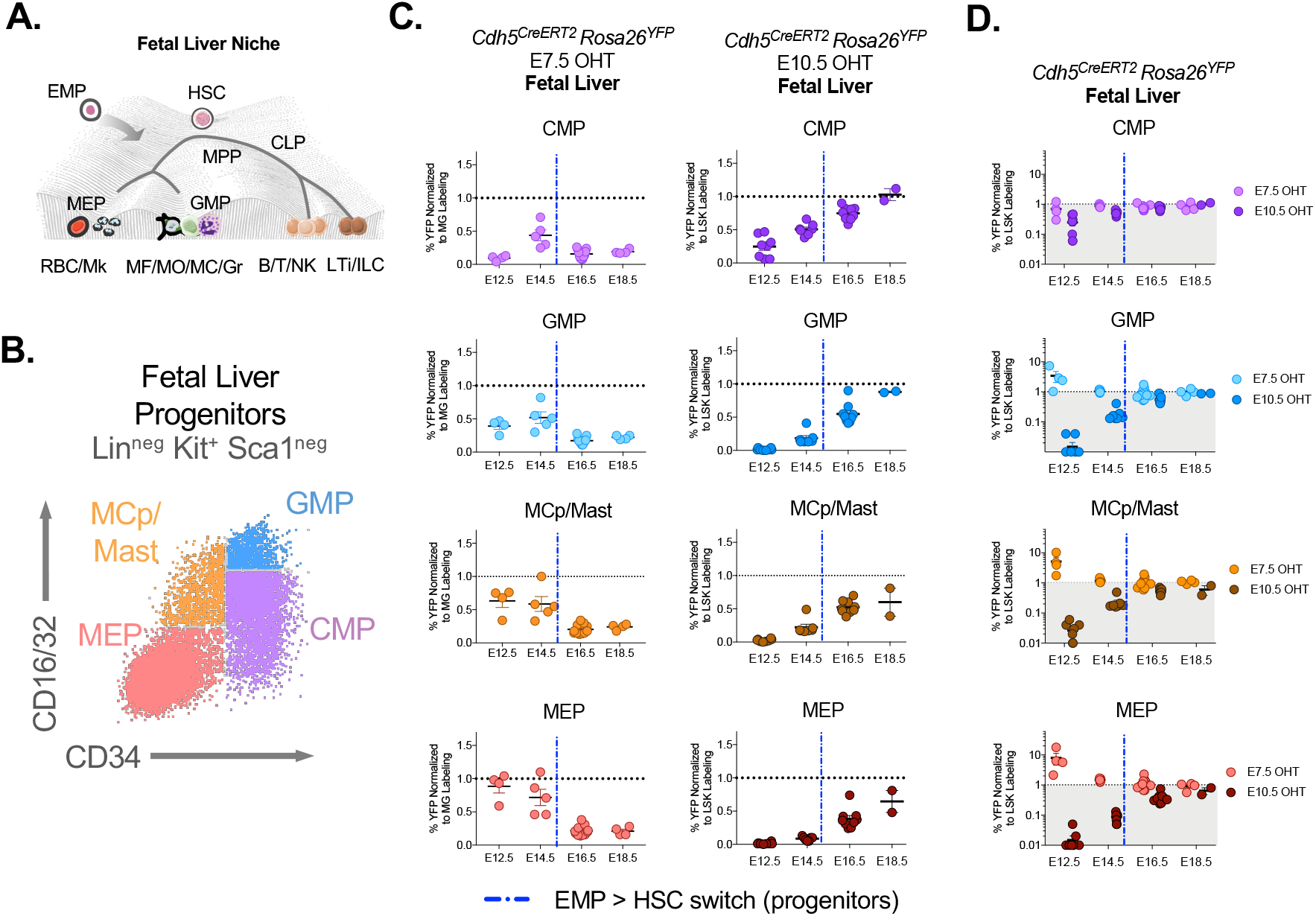
Delayed allocation of fetal HSC to erythroid and myeloid committed progenitor compartments. **A.** Simplified hematopoieteic progenitor hierarchy superimposed on a Waddington landscape to illustrate the bifurcated cascade of HSC into common myeloid progenitors (CMP) and common lymphoid progenitors (CLP) downstream of multipotent progenitors (MPP). CMP further give rise to megakaryocyte-erythrocyte progenitors (MEP) and granulocyte-monocyte progenitors (GMP). Hematopoiesis in the fetal liver niche is further complicated by the additional layer of EMP that migrate from the yolk sac. **B.** Gating of CMP, GMP, MEP and MCp/Mast (example from E12.5 fetal liver) progenitor subsets based on cell surface intensity of CD16/32 and CD34 among LK (Lin^neg^ Kit^+^ Sca1^neg^). **C.** Dynamics of EMP and HSC derived hematopoietic progenitor subsets labeled by *Cdh5*^*CreERT2*^ *Rosa26*^*YFP*^ with E7.5 OHT (left) or E10.5 OHT (right). The percent of YFP labeling among progenitor subsets was normalized to percent YFP among microglia (CD45^+^ F4/80^+^) for E7.5 OHT or LSK (Lin^neg^ Sca1^+^ Kit^+^) for E10.5 OHT (see Figure S1A). **D.** Representation of data from Panel C with normalization of both injection timepoints to LSK labeling. MG, microglia; RBC, red blood cell; Mk, megakaryocyte; MCp, mast cell progenitor; MC, mast cell; MO, monocyte; Nt, neutrophil; MF, macrophage; LTi, lymphoid tissue inducer; ILC, innate lymphoid cell. Data are represented as mean ± SEM.

Using the *Cdh5*^*CreERT2*^ *Rosa26*^*YFP*^ pulse chase labeling approach, the relative contributions of EMP or HSC were compared among CMP, GMP, MCp/Mast and MEP populations (**Figure 2C**, **Figure S2C**). The same normalization approach was used as described for *Cdh5*^*CreERT2*^ *Rosa26*^*YFP*^ labeling of mature cells (**Figure 1A,C**). Erythroid (MEP) and myeloid (GMP and MCp/Mast) progenitor compartments were populated between E12.5 and E15.5 by EMP progeny (labeled by E7.5 OHT). Conversely, CMP appeared to be largely representative of HSC progeny in the fetal liver (labeled by E10.5 OHT). This was better visualized by normalizing YFP^+^ progenitors to YFP^+^ LSK (**Figure 2D**), since both injection timepoints label LSK. With this representation, it can be seen that the EMP wave never exceeded equilibration with CMP from E12.5 onwards because these CMP were almost exclusively from the HSC wave. HSC progeny began allocating to erythroid and myeloid compartments from around E15 onwards, one day prior to the EMP-to-HSC switch observed for mature cells. The numbers of HSC progeny per fetal liver (**Figure S2C**) peaked at E16.5 followed by a decline, possibly due to transition in the hematopoietic niche to the bone marrow that occurs prior to birth.

### Early amplification erythro-myeloid progenitor intermediates by EMP

Further study of the early expansion and commitment of EMP erythroid and myeloid progenitors using the *Cdh5*^*CreERT2*^ *Rosa26*^*YFP*^ approach was complicated by the concomitant labeling of EMP and LSK with E7.5 OHT injection. Instead, we used the *Csf1r*^*MeriCreMer*^ *Rosa26*^*YFP*^ model that reliably marks newly emerged EMP independently of HSC (**Figure 3, Figure S3**). Since EMP are produced in the yolk sac successively over the course of several days, we assessed labeling by injection of OHT at E8.5 or E9.5, however later timepoints were not pursued as they result in labeling of HSC (Elsaid et al., 2021). Fetal liver LK (Lin^neg^ Kit^+^ Sca1^neg^) were collected between E10.5 and E14.5 and characterized by flow cytometry.

**Figure 3.**
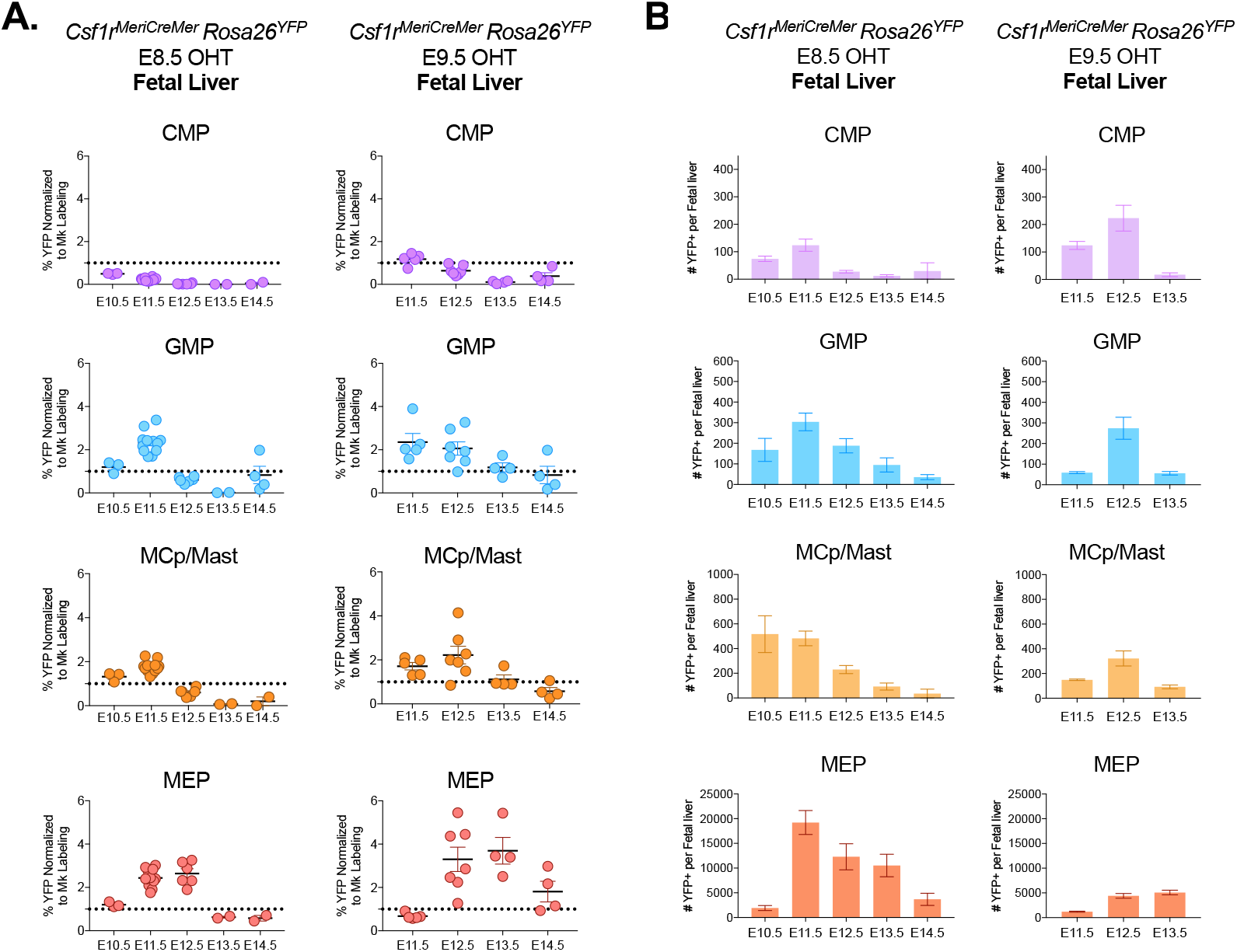
Transit Amplification of EMP-Derived Erythroid and Myeloid Progenitor Subsets in the Early Fetal Liver. **A.** Dynamics of yolk sac derived EMP labeled by *Csf1r*^*MeriCreMer*^ *Rosa26*^*YFP*^ with E8.5 OHT are represented by percent YFP normalized to labeling among megakaryocytes (Mk, Kit^neg^ CD45^neg^ CD41^hi^). EMP contribute poorly to CMP, but undergo expansion as GMP, MEP or MCp (mast cell progenitor) intermediates. **B.** Numbers of EMP derived CMP, GMP, MCp/Mast and MEP labeled by *Csf1r*^*MeriCreMer*^ *Rosa26*^*YFP*^ with E8.5 OHT or E9.5 OHT in the fetal liver. The number of YFP^+^ fetal liver hematopoietic progenitors peak 72 hours after injection regardless of injection timepoint. Data are represented as mean ± SEM.

For EMP progeny that were labeled by E8.5 and E9.5 OHT injection, the percentage of YFP labeling among all hematopoietic progenitors (LK) was the same when comparing LK in the yolk sac to those in the peripheral blood. However, the labeling of LK in the fetal liver remained elevated at E11.5 (for E8.5 pulsed) and E12.5 (for E9.5 pulsed) (**Figure S3A**) indicating an expansion of EMP-derived LK in the fetal liver during these times.

Differences in labeling efficiency of EMP by E9.5 compared to E8.5 OHT injection were observed, perhaps explained by an abundance of EMP-derived erythroid committed progenitors generated in the yolk sac between E9.5 and E10.5 (Iturri et al., 2021) that are no longer expressing *Csf1r*. There was also higher efficiency of microglia labeling likely due to labeling of pre-macrophages expressing *Csf1r* at E9.5. Therefore, for comparability of E8.5 and E9.5 pulse chase labeled progenitor cells (LK, Lin^neg^ Kit^+^), we could not normalize the percent labeling among LK subsets to labeling of microglia, but rather we normalized to labeling of megakaryocytes (Mk, Kit^neg^ CD45^neg^ CD41^hi^) (**Figure 3A**), since Mk at these stages are yolk sac derived (Iturri et al., 2021). There was limited contribution of EMP to the CMP compartment with either E8.5 or E9.5 OHT injection, whereas EMP-derived GMP, MCp/Mast and MEP expanded at E11.5 and E12.5 respectively (**Figure 3A-B**). EMP progeny that were pulse chase labeled at E8.5 or E9.5 generated similar numbers of YFP^+^ myeloid progenitors within the fetal liver when quantified 72 hours after the time of OHT injection.

Therefore, erythroid- and myeloid-committed progenitors in the early fetal liver are rapidly generated by EMP. This is followed by late and gradual commitment from HSC-derived CMP, suggesting that fetal HSC may undergo tighter regulation of erythro-myeloid differentiation compared to EMP or that they may compete for signals from the niche. Early EMP-derived myeloid progenitors (GMP and MCp/Mast) from two different timepoints undergo 72 hours of transit amplification prior to differentiation, linking their time of birth in the yolk sac to their time of differentiation, suggesting that this may be intrinsically regulated.

### Three pathways of fetal liver EMP myelopoiesis

We observed that the production of mature myeloid cells during development is ontogeny- and lineage-specific. This led us to further examine the molecular basis for myeloid progenitor heterogeneity in the fetal liver using single-cell RNA sequencing (scRNA-seq). Since the ontogeny of erythro-myeloid progenitors in the fetal liver prior to E15 could be traced back to yolk sac EMP, we used *Csf1r*^*MeriCreMer*^ *Rosa26*^*YFP*^ embryos pulsed at E8.5 by OHT injection to isolate EMP-derived LK (Lin^neg^ Kit^+^ YFP^+^) from E10.5 and E12.5 fetal livers.

We performed massively parallel single-cell RNA sequencing (MARS-seq) (Jaitin et al., 2014; Keren-Shaul et al., 2019) because it permits index sorting for retrospective analysis of cell surface markers corresponding to individual cell transcriptomes. 1,590 cells from E10.5 fetal liver and 1,405 cells and E12.5 were analyzed (**Figure S4A-C**). Due to the overwhelming abundance of erythroid progenitors in the fetal liver, sampling of E12.5 fetal liver MEP was selectively downsized to enrich for non-MEP (see Methods). UMAP representation of clustered cells demonstrated two major pan-myeloid (clusters 2-5) and pan-erythroid (clusters 6-10) branches (**Figure 4A**) exemplified by up- or down-regulation of the erythroid and myeloid specific transcription factors PU1 (*Sfpi1*) and GATA1 (*Gata1*) (**Figure S4D**).

**Figure 4.**
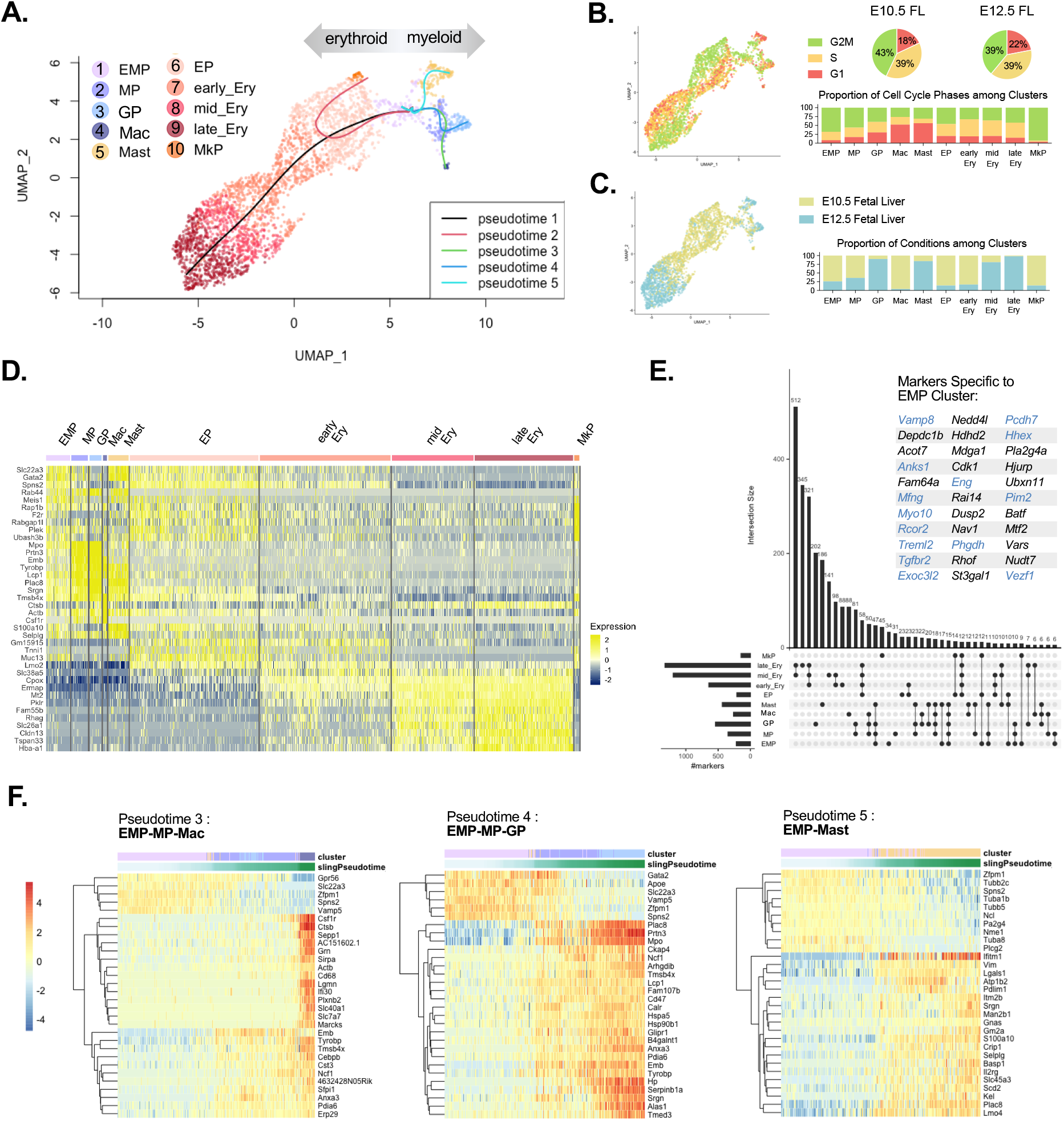
Molecular Heterogeneity among Fetal Liver EMP-derived Progenitors. **A.** UMAP representatiion of Lin^neg^ Kit^+^ YFP^+^ fetal liver cells isolated from *Csf1r*^*MeriCreMer*^ *Rosa26*^*YFP*^ embryos pulsed with E8.5 OHT (1,590 cells from E10.5 and 1,405 cells from E12.5, see Figure S3A-C for strategy and filtering parameters). Cells from two conditions (E10.5 fetal liver and E12.5 fetal liver) were merged using fastMNN and clustered using a walktrap algorithm. Pseudotime was inferred using Slingshot. Cells spread across two major myeloid (clusters 2-5) and erythroid (clusters 6-10) branches. **B.** UMAP colored by cell cycle phase. Proportion of cell cycle phase assignments per cluster (bar graph) or by condition (pie charts). **C.** UMAP colored by condition. Proportion of conditions per cluster (bar graph). **D.** The heatmap shows top 5 selected cluster markers (identified as differentially expressed in at least half of pairwise comparisons). Notably, the EMP cluster was characterized by mixed expression of both myeloid and erythroid genes. **E.** UpSet plot illustrating shared markers between clusters. Horizontal bars show the total number of markers per cluster. Vertical bars represent the size of the corresponding interaction represented in the dotplot. There were 34 markers specific to the EMP cluster (endothelial cell markers in blue). **F.** Heatmaps of differential gene expression along inferred pseudotime lineages from the EMP cluster to myeloid cell types (trajectories indicated in Panel A). EMP, erythromyeloid progenitor; MP, myeloid progenitor; GP, granulocyte progenitor; Mac, macrophage; EP, erythroid progenitor; Ery, erythroid; MkP megakaryocyte progenitor.

Clustering was not driven by cell cycle phase (**Figure 4B**) or differences in embryonic stage (**Figure 4C**), but rather by expression of cluster specific markers (**Figure 4D**). Erythroid and myeloid branches were bridged by the EMP cluster that was characterized by combinatorial erythroid and myeloid genes and limited expression of markers that were not shared among other clusters (**Figure 4E**). To assign a cell type to each cluster, we performed pairwise correlation using the average cluster expression profiles from our dataset compared to those of previously annotated E14 fetal liver hematopoietic cells from the Mouse Cell Atlas project (Han et al., 2018) (**Figure S4E**). The EMP cluster correlated most closely with stem/progenitor cells while additional myeloid clusters were identified as GP (granulocyte progenitors, highly correlated with Elane^hi^ neutrophils and less with more mature Ngp^hi^ neutrophils) and Mast cells.

Pseudotime analysis traced pathways extending to three budding branches of myeloid progenitor differentiation (**Figure 4F**). Most represented among these were the clusters of Mast and GP whereas the numbers of progenitors going towards macrophage fate were not well defined. This was evident by the poorly delineated trajectory leading towards a small number of pre-macrophages (cluster 4, Mac), identified as expressing F4/80 (*Adgre1/Emr1*) that were incidentally captured during the sorting process. This supports the notion that tissue macrophages differentiate in the yolk sac and traffic throughout the embryo from E10.5-E12.5 (Stremmel et al., 2018) where they can self-propagate within target tissues.

### Lineage priming of EMP-derived progenitors in the fetal liver

We observed that EMP in the fetal liver preferentially produce mature mast and neutrophils *in vivo* and that the progenitors to these mature cells expand from E11.5-E12.5 within immunophenotypically defined GMP and MCp/Mast populations. scRNA-Seq revealed molecular signatures of EMP-derived myeloid progenitors in the fetal liver that were specifically enriched for mast and granulocyte potential, but lacking in monocyte/macrophage progenitor identity. These differential preferences for myeloid lineage output from EMP versus HSC raised the questions of whether EMP myeloid progenitors utilize the same pathways or can be characterized by the same immunophenotypes as those used to study HSC.

To examine cell intrinsic priming of EMP-derived myeloid progenitors in the fetal liver, we selected key transcription factors (TF) involved in lineage priming of adult myeloid progenitors (Paul et al., 2015) and examined their average expression levels among each cluster from our scRNA-seq dataset (**Figure 5A**). Erythroid priming TF (*Phf10* and *Klf1* for red blood cells, *Fli1* and *Pbx1* for megakaryocytes) were highly conserved. Expression of *Irf8* was mainly within the Mac cluster (with high levels of *Maf*) along with lower expression in the MP cluster. Expression of genes identifying monocytes or monocyte progenitors (*Ly6c2, Flt3, Ccr2, S100a4*) were mostly undetectable (**Figure S5A**) thereby showing minimal monocyte lineage priming among EMP progeny in the fetal liver. GP expressed high levels of the neutrophil priming TF *Cebpe* along with low level of *Gfi1*. The TF Lmo4, identified as basophilic among adult myeloid progenitors, was very highly expressed in the Mast cluster along with other TF identified by our analysis to be specifically enriched among this cluster (*Foxo3, Gata2, Sox4*).

**Figure 5.**
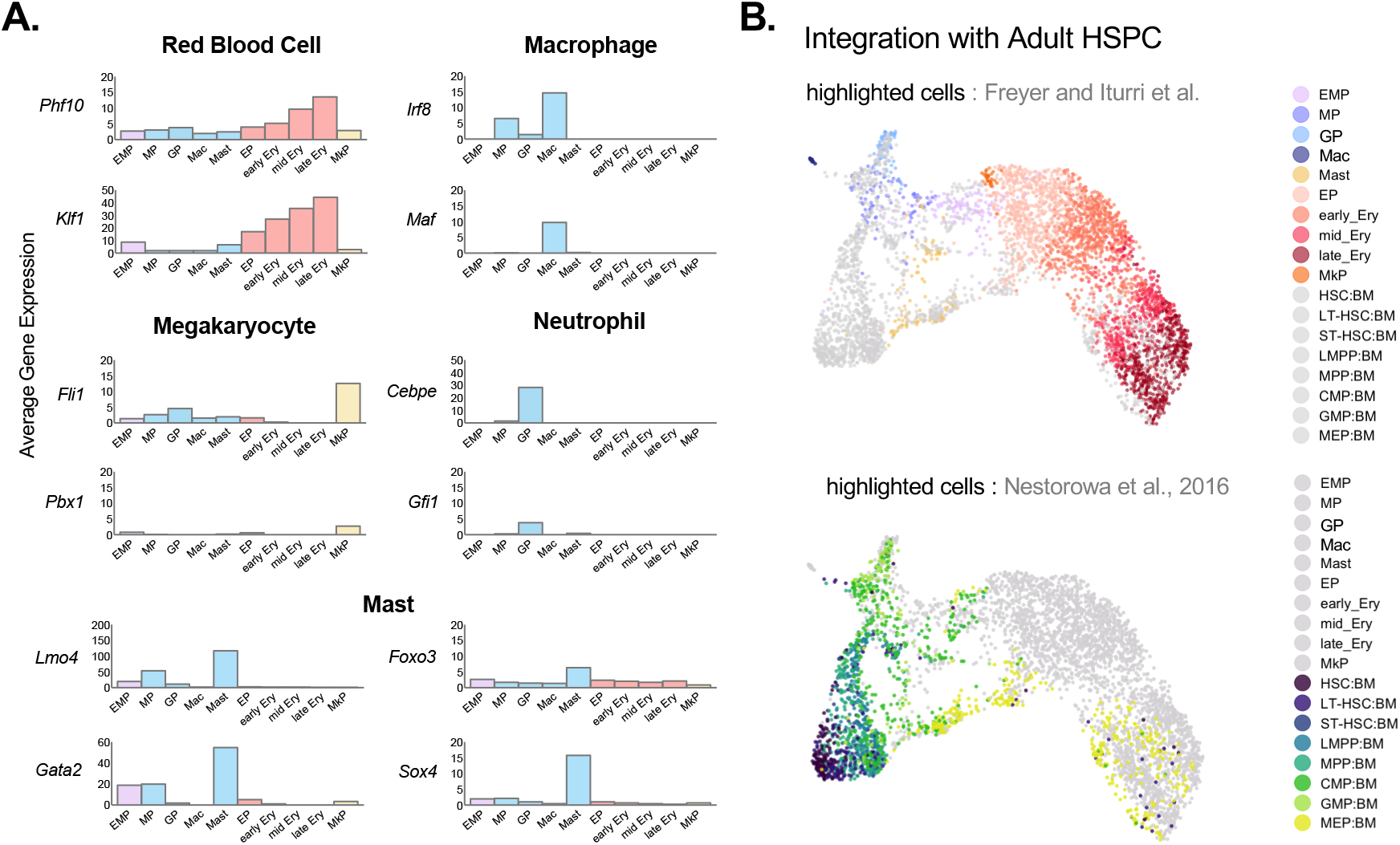
Transcriptional Priming of EMP-derived Progenitors in the Fetal Liver. **A.** Average expression of lineage defining transcription factors by cluster. Transcription factors were selected based on previously described lineage priming of adult myeloid progenitors (Paul et al., 2015). **B.** UMAP representation of fetal liver EMP-derived progenitors integrated with previously published adult HSC/HSPC from Nestorowa et al., 2016. Nestorowa cluster names were based on indexed FACS analysis and therefore represent cell identity according immunophenotyping and not by molecular signature. For visualization of individual clusters, see Figure S5A.

To determine the molecular relationship between EMP-derived myeloid progenitors in the fetal liver and HSC-derived myeloid progenitors from the bone marrow that have been isolated by immunophenotyping (CMP, GMP, MEP), we integrated our dataset with published Smart-seq2 analysis of index-sorted HSPC from adult bone marrow (**Figure 5B**, **Figure S5B**) (Nestorowa et al., 2016). Considering the failure of EMP to demonstrate lymphoid potential (Elsaid et al., 2021), it was expected that EMP progeny did not integrate with adult long-term HSC (LT-HSC) or short-term HSC (ST-HSC), nor with HSPC upstream of CMP such as multipotent progenitors (MPP) or lymphoid multipotent progenitors (LMPP). However, adult bone marrow GMP and a subset of CMP locally mapped to regions of the UMAP populated by cells from our EMP, MP and GP clusters. Adult bone marrow MEP followed the same trajectory as EMP-derived MEP although the cells did not integrate until more advanced stages of erythroid differentiation (late_Ery). This may be explained by differences in fetal versus adult erythropoiesis, for example the rapid loss of CD45 expression on fetal red blood progenitors (Soares-da-Silva et al., 2021).

### Single-cell heterogeneity of EMP-derived myeloid progenitor immunophenotypes

TF and integration analyses showed that EMP-derived myeloid progenitors in the fetal liver share molecular characteristics with adult bone marrow HSC-derived CMP, GMP and MEP. To determine whether the isolation of EMP-derived myeloid progenitors from the fetal liver by immunophenotyping (CMP, GMP, MEP gating by flow cytometry) is predictive of their lineage potential, we generated dot plots of the cells from our scRNA-seq according to the median fluorescence intensity (MFI) of cell surface CD16/23 and CD34 captured by index sorting. Then we overlaid cells from each cluster onto the two-dimensional plots (**Figure 6A-B**, **Figure S6**). EMP, represented mainly by E10.5 cells, were localized mostly in the CMP region then more frequently among CD16/32^hi^ CD34^lo^ cells at E12.5. Other myeloid progenitors (MP and GP clusters) were also mostly within the CMP gate at E10.5, then later found predominantly in the GP gate at E12.5. Manual gating of MEP was efficient at including the vast majority of erythroid progenitors (EP, early_Ery, mid_Ery and late_Ery clusters, **Figure S6**) whereas this was largely not predictive for megakaryocyte progenitors (MkP cluster), as has been previously reported for megakaryocyte progenitors in the adult bone marrow (Paul et al., 2015).

**Figure 6.**
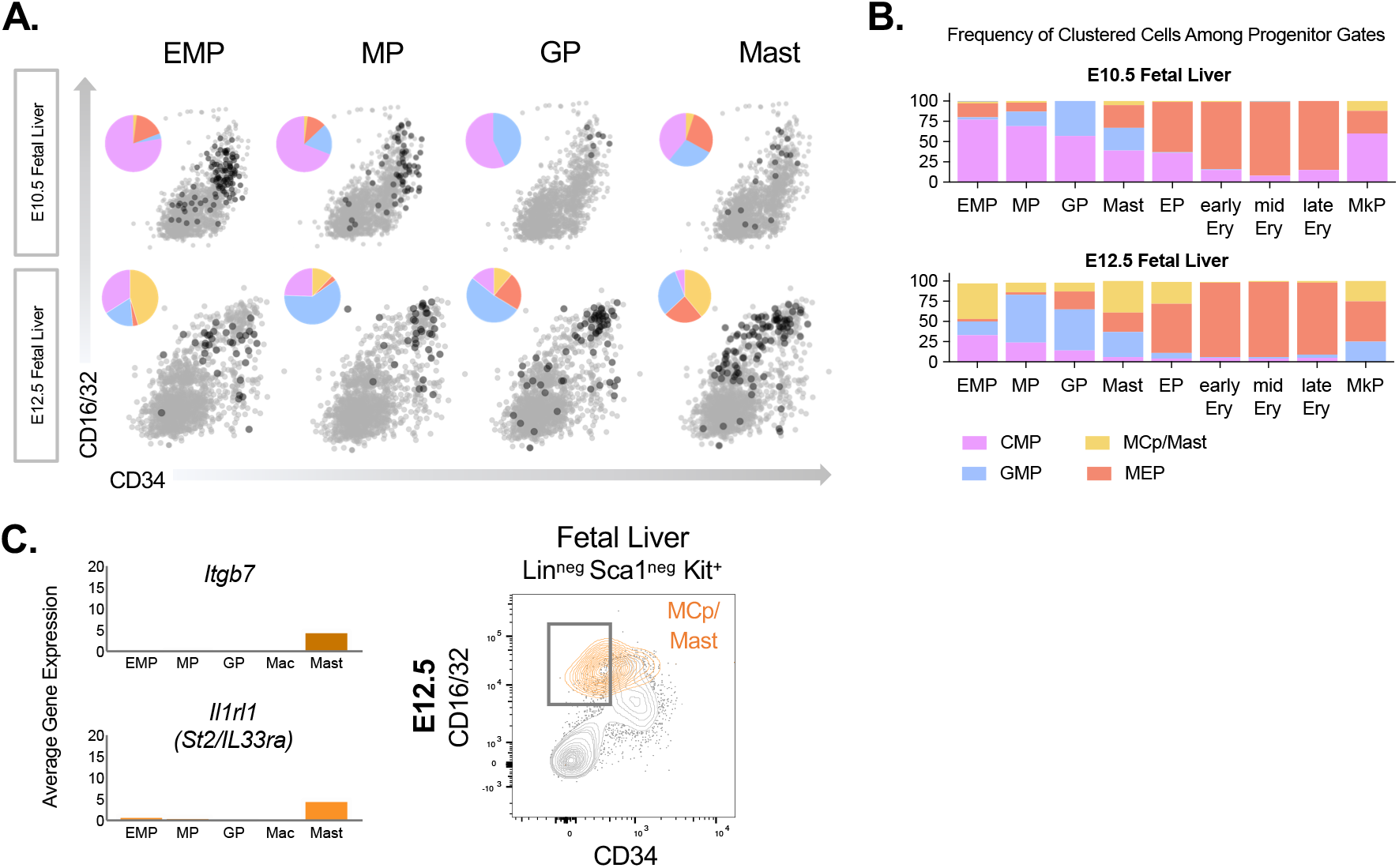
Single cell immunophenotype of EMP-derived Progenitors in the Fetal Liver. **A.** Index sorting classification of EMP progeny from scRNASeq data using conventional CD16/32 versus CD34 gating to define CMP, GMP, MCp/Mast and MEP progenitor subsets for retrospective plotting of individually sequenced cells. **B.** Quantification of the relative number of cells per cluster that lie within each of the traditional gates. For index sorting classification of Mac and Ery clusters, see Figure S6B. **C.** Genes expressing cell surface markers that were specific to the Mast cluster (left). Flow cytometry of E12.5 fetal liver LK (right) with overlay of ITGB7^+^ LK (orange).

Notably, the Mast cluster was strongly represented among CD16/32^hi^ CD34^lo^ cells demonstrating that non-conventional gating may enrich for mast cell progenitors in the fetal liver. Therefore, we searched for genes that met the criteria of being specifically expressed in the Mast cluster and that encode cell surface proteins for which there are commercially available fluorescently-coupled antibodies. We identified *Itgb7* and *Il1rl1 (ST2/Il33ra)* (**Figure 6C**) as candidates and observed some fetal liver LK that expressed cell surface ITGB7 at E12.5. As expected based on the scRNA-seq with index sorting, flow cytometry demonstrated that these ITGB7^+^ LK were present among GMP but more enriched among CD16/32^hi^ CD34^lo^ cells (**Figure 6C**). Therefore, immunophenotypic GMP in the early fetal liver are EMP-derived and composed of a heterogeneous mix of cells that are molecularly primed for production of neutrophils and mast cells.

## DISCUSSION

During development, the fetal liver is the prime hematopoietic organ, supporting differentiation from two unique pools of hematopoietic progenitors with distinct extra-embryonic (EMP) versus intra-embryonic (HSC) origins. We used two independent genetic pulse chase labeling models to lineage trace the origins of mature cells from EMP or HSC.

Surprisingly, the contribution of HSC to mature myeloid cells in the fetal liver was only evident late in gestation (>E16). Meanwhile, EMP robustly generated mast cells as well as neutrophils that were still found in circulation after birth (data not shown). The significant role of EMP in generating myeloid cells during development is similar to what has been reported for mature erythroid cells (Soares-da-Silva et al., 2021). On the contrary, HSC are an early source of lymphoid cells that cannot be generated by EMP (Elsaid et al., 2021) and which may be partially replaced later in life by adult HSC. Collectively, our findings have led us to propose a model in which EMP and HSC ‘divide and conquer’ to build a functional hematopoietic system during development that serves to sustain survival and growth of the embryo while also laying the foundations for a complex layered immune system that is maintained throughout adult life. This division of labor may be a mechanism for managing the increased demands that are unique to the gestational period.

Our study has pinpointed an ‘EMP-to-HSC-switch’ that occurs around E16 in fetal liver hematopoiesis, demonstrated by ontogeny-specific production of mature cells and reflected at the progenitor level one day earlier. Investigation into the allocation of EMP and HSC to transit amplifying progenitor subsets showed that EMP rapidly generate GMP, MCp and MEP while HSC lag behind in this process. The amplification of MEP-derived intermediates occurred 72 hours after their labeling in the yolk sac, regardless of the injection timepoint. This could possibly reflect intrinsic programming or alternatively may indicate the time needed for exposure to factors in the fetal liver niche. On the other hand, HSC were the primary source of phenotypic CMP and only contributed to GMP, MCp and MEP intermediates late in gestation. It remains unknown how these differences in timing are regulated. Addressing this question is complicated by the timing of colonization of the fetal liver niche by EMP and HSC that occurs early, is separated only by 24 hours and is a continuous process that spans several days of EMP and HSC production in their primary niches. Furthermore, the relationship between hematopoietic colonization and maturation of the fetal liver niche would need to be taken into consideration. One possibility is that EMP are more sensitized to growth factors in the niche and therefore outcompete their HSC counterparts. Since EMP do not appear to self-renew in the fetal liver, their depletion with time would then allow for HSC to thrive in their absence.

The predominance of EMP-derived myeloid cells in the early fetal liver led us to further investigate the lineage- and ontogeny-specific characteristics of EMP differentiation at the molecular level. scRNA-seq identified three pathways to myeloid cell differentiation from EMP. The pathways to producing mast cells and neutrophils were dominant while there were few progenitors linked to differentiation of macrophages. We attributed this to the early differentiation of macrophages in the yolk sac niche. EMP-derived myeloid progenitors expressed key lineage defining transcription factors showing that many had already been primed for lineage choice. We did not identify gene signatures for monocyte-like progenitors, in line with our *in vivo* pulse chase labeling results.

Index sorting exposed the molecular heterogeneity of EMP-derived CMP, GMP, MCp and MEP (that were classified by cell surface intensity of CD16/32 and CD34) and demonstrated that conventional gating approaches were not sufficient to isolate lineage primed cells. For example, putative MCp were identified as ITGB7^+^ cells mainly localized in the CD16/32^hi^ CD34^lo^ region but also overlapping with GMP. It will interesting to validate the potential of putative MCp by *in vitro* colony-forming unit (CFU) assays and to test the enrichment of MCp from the fetal liver using a modified gating approach with ITGB7.

Other studies have proposed an EMP origin of monocytes that pass through a monocyte-dendritic progenitor (MDP) to cMoP (common monocyte progenitor) pathway (Hoeffel et al., 2015). However, our *in vivo* lineage tracing did not provide evidence for a significant contribution of EMP to fetal liver monocytes. Instead, it appeared that HSC were more likely than EMP to make fetal liver monocytes and that this may largely be regulated by a pathway that was uncoupled to the production of neutrophils, for example via the MDP-cMoP-MO pathway. MDP are defined as CSF1R^+^ Kit^+^ Flt3^+^ and expression of Flt3 is localized to the HSC-derived CMP compartment of the fetal liver (data not shown). EMP-derived progenitors did not express Flt3 thus making it likely that fetal monocytes generated from MDP intermediates are HSC-derived. However, we cannot exclude the possibility that EMP produce some monocytes through a GMP intermediate (Yáñez et al., 2017).

It remains unclear whether fetal liver GMP (derived from either EMP or HSC) are capable to generate monocytes/macrophages *in vivo.* Therefore it is of great interest to isolate EMP-derived versus HSC-derived GMP to first test their differentiation capacity *in vitro* by CFU assay. It is possible that saturating levels of cytokines will permit production of mixed granulocyte-monocyte/macrophage colonies regardless of ontogeny. This would suggest that lineage decisions made *in vivo* are driven by the microenvironment. In that case it would be essential to perform adoptive transfer *in utero* or in neonates to compare the potential of isolated progenitors within the liver niche.

## METHODS

### Mice

All mice used in this study have been previously described. Experimental procedures, housing and husbandry were conducted in compliance with the regulatory guidelines of the Institut Pasteur Committee for Ethics and Animal Experimentation (CETEA, dap160091). Strains included *Csf1r*^*MeriCreMer*^ transgenic (FVB background, MGI:5433683), *Rosa26*^*YFP*^ knock-in (C57Bl6 background, MGI:2449038) and *Cdh5*^*CreERT2*^ transgenic (C57Bl6 background, MGI:3848982) mice. To generate embryos for pulse chase labeling experiments, *Rosa26*^*YFP/YFP*^ males were crossed to *Csf1r*^*MeriCreMer*/+^ or *Cdh5*^*CreERT2*/+^ females in timed matings and the date of vaginal plug was considered to be embryonic day E0.5. Embryonic stage was validated by counting somite pairs when possible (E10.5-11.5) or using morphological landmarks.

### PCR Genotyping

DNA from tissue biopsy or embryonic sample was extracted using the Hot Shot method : tissue was lysed in 25μL (embryonic tissue) or 50μL (postnatal tissue) of alkaline lysis solution (25mM NaOH, 0.2mM EDTA) at 95°C for 20 minutes then equal amount of neutralizing solution was added (40mM Tris-HCl). PCR was performed using a BIOTAQ kit (Bioline). *Csf1r*^*MeriCreMer*^ mice were genotyped using iCre-F-YL (5’- GTCTCCAACCTGCTGACTGTGC-3’) and iCre-R (5’-CCAATGCTGTGTCCCTGGTGATG-3’) oligos with 65°C annealing temperature (400bp product). *Rosa26*^*YFP*^ mice were genotyped using R_YFP_1 (5’- AAGTCGCTCTGAGTTGTTAT-3’), R_YFP_2 (5’- GGAGCGGGAGAAATGGATATG-3’) and R_YFP_3 (5’- GCGAAGAGTTTGTCCTCAACC-3’) oligos with 60°C annealing temperature (525bp product for wild-type and 300bp product for knock-in). *Cdh5*^*CreERT2*^ mice were genotyped using Cre_Fwd (5’- GCCGAAATTGCCAGAATCAG-3’) and Cre_Rev (5’-ACATTGGTCCAGCCACCAGC-3’) oligos with 60°C annealing temperature (430bp product).

### 4-Hydroxytamoxifen (OHT) Preparation and Injection

50mg/mL stocks of 4-hydroxytamoxifen (OHT) (Sigma, H7904-25MG) were prepared by resuspending 25mg of OHT in 250μL of ethanol by vortexing for 15 minutes then sonicating for 30 minutes, followed by adding 250μL of Kolliphor (Sigma C5135-500G) and sonicating again for 30 minutes. 60μL aliquots (3mg) were stored at −20°C for up to 6 months. 10mg/mL stocks of progesterone (P3972-5G) were prepared by resuspending 25 mg of progesterone in 250μL of ethanol, vortexing, adding 2250μL of sunflower oil (Sigma S5007-250ML) and storing at −4°C for up to 2 weeks. For pulse labeling, OHT was co-injected with progesterone to reduce the risk of abortion. An injection solution was prepared by thawing 60μL of 50mg/mL OHT and sonicating for 30 minutes, then adding 240μL of 0.9% NaCl and sonicating for 30 minutes, then adding 150μL of 10mg/mL progesterone and sonicating for 30 minutes. For pulse labeling using the *Csf1r*^*MeriCreMer*^ strain, females were weighed on day 8 of pregnancy and injected with 11μL/g for final concentration of 75μg/g OHT and 37.5μg/g progesterone. For pulse labeling using the *Cdh5*^*CreERT2*^ strain, females were weighed on day 7 of pregnancy and injected with 8μL/g for final concentration of 50μg/g OHT and 25μg/g progesterone. Injections were performed using 1mL syringes with 29G needles at 13h. Live births of pulse chase labeled mice were complicated by OHT injection during pregnancy, therefore we performed caesarean sectioning followed by fostering with FVB females that had given birth within less than 5 days.

### Flow Cytometry

For embryonic experiments, the protocol for preparing cells for flow cytometry analysis has been previously described in greater detail (Iturri et al., 2017). The pregnant mouse was killed by cervical dislocation and the uterus was removed and placed in PBS. Embryos were isolated by opening the uterus and removing the extraembryonic membranes without severing the vitelline and umbilical vessels. To collect peripheral blood, the embryo was separated from the placenta by severing the vitelline and umbilical vessels and rapidly placing the embryo in a 12-well plate with 2mL of 2mM EDTA. Bleeding was facilitated by completely detaching the vitelline and umbilical vessels from the embryo and leaving them for 10 minutes. Organs were dissected in cold PBS and placed in 24-well plates with 500μL cold digestion buffer composed of PBS with 1mg/mL collagenase D (Sigma 11088882001), 100 U/mL DNaseI (DN25-100mg) and 3% Fetal Bovine Serum. When dissection was complete, the digestion plates were placed at 37°C for 30 minutes. Digestion was stopped by transferring tissue to 100μm strainers placed in 6-well plates with 5mL filtered FACS Buffer (0.5% BSA and 2mM EDTA in PBS), then the cells were passed through the strainer by mashing with the piston of a 5mL syringe. Cells were pelleted at 320*g* for 7 minutes. Blocking was performed with 5% FBS and 1:20 Mouse IgG (Interchim 015-000-003) in FACS Buffer followed by 30 minutes of antibody staining. Cells were washed and incubated with fluorescently-conjugated streptavidin for 20 minutes.

For experiments with adult mice, pieces of organs were patted dry and weighed prior to digestion and 2.4mg Dispase was added to the digestion buffer. Tissue was minced with scissors before starting digestion. Blood was collected by retro-orbital bleeding. Red blood cell lysis was performed on all organs one time and twice for blood. Bone marrow was flushed from the femur and tibia using a 25G needle. Stained cells were passed on the BD Symphony A5 cytometer (Diva software).

To quantify cells, the following formula was used : (# of cells acquired) x (volume of resuspended cells after staining and washing/volume of cells acquired) x (volume of cell suspension in blocking buffer prior to staining/volume of cells plated for staining). The Symphony does not record volume, so the volume of cells acquired was noted after passing an entire aliquot. Results were analyzed and plots generated using FlowJo.

### Antibodies

| Antigen | Conjugates | Clone |
| --- | --- | --- |
| CD16/32 | BUV737 | 2.4G2 |
| CD45 | BUV661, BUV395 | 30-F11 |
| CD45.2 | APC-Cy7 | 104 |
| CD41 | BUV395 | MWReg30 |
| F4/80 | BV421, BV785, biotin | BM8 |
| Ly6G | BV711 | 1A8 |
| Ly6C | BV605 | HK1.4 |
| Sca1 | BV510 | D7 |
| CD34 | eF450 | RAM34 |
| Itgb7 | PE-Cy7 | FIB504 |
| Ter119 | biotin | TER-119 |
| CD19 | biotin | 1D3 |
| CD3e | biotin | 145-2C11 |
| Gr1 | biotin | RB6-8C5 |
| NK1.1 | biotin | PK136 |
| CD115 | APC, PE-CF594 | AFS98 |
| Flt3 | PE, PerCP-eF710 | A2F10 |
| Kit | APC-Cy7, PE | 2B8 |
| CD11b | AF700, PE-Cy7 | M1/70 |
| AA4.1 | APC, PE-CF594 | AA4.1 |
| CD61 | PE-Cy7 | 2C9.G2 |
| CD71 | AlexaFluor647 | C2 |
| CD150 | PE-Cy7 | mShad150 |
| CD48 | APC | HM48-1 |
| SAV | PE-Cy5, BUV737 |  |

### Single-Cell Library Preparation

Cells were prepared for fluorescence-activated cell sorting (FACS) as described for flow cytometry. Embryos were screened for YFP expression by rapidly dissociating the head with a syringe, filtering, then passing unstained cells on the Beckman Coulter Cytoflex cytometer. YFP^+^ embryos were pooled. Lineage cells (Ter119^+^ F4/80^+^ Gr1^+^) were depleted using magnetic anti-biotin Microbeads (1:5) (Miltenyi Biotec 130-090-485) and MS columns (Miltenyi Biotec 130-042-201). Lin^neg^ Kit^+^ YFP^+^ cells were index sorted using a 70μm nozzle onto 384-well capture plates (10% Triton X-100, 40U/μL RNasin plus, dispensed at 4°C using the Bravo pipetting robot) with 4 wells that were left empty as control. Since MEP represent more than 90% of fetal liver LK at this E12.5, the sorting of E12.5 MEP were limited to half of each plate in order to enrich for myeloid progenitors (this selection was not made for sorting of E10.5 fetal liver). Plates were stored at −80°C until the library preparation step. During library preparation, cDNA from each 384-well plate was divided into two pools termed ‘A’ and ‘H’ half-plates which each contained cDNA from 192 uniquely barcoded wells (barcode oligos prepared by Baptiste Saudemont, UTECHS CB). The protocol and reagents for MARS-Seq library preparation have already been described in detail (Jaitin et al., 2014; Keren-Shaul et al., 2019).

### High-Throughput Sequencing

Libraries were pooled in equimolar amounts of 20fmol per half-plate for final concentration of 1-2nM. Pooled libraries were sequenced using the NextSeq 500/550 High Output Kit v2 for 75 cycles (Illumina FC-404-2005). Base calling accuracy was greater than 80% (>=Q30, >40G) and 75-80% clusters passed filter. Analyzed data included results from two different sequencing runs. Barcode extraction, alignment, generation of QC reports and creation of umi.tab expression tables were performed by Yann Loe Mie (Bioinformatics and Biostatistics Hub, pipeline available at http://compgenomics.weizmann.ac.il/tanay/?page_id=672). Raw and processed data will be made available on GEO.

### Cell and Gene Filtering

The number of UMIs (counts) and features (detected genes) were filtered to eliminate cells with less than 3,000 detected genes. Cells were filtered to remove those that had a mitochondrial gene fraction that was greater than 5%. Median Fluorescence Intensity (MFI) values for CD34 and CD16/32 from index sorting were matched to individual transcriptomes by their well ID and added to metadata of a Seurat object. Doublet detection for each plate was performed using default parameters of scDblFinder then filtered to remove cells classified as doublet. Expressed genes were defined as being detected in at least 5% of cells from each condition (E10.5_Fetal_Liver, E12.5_Fetal_Liver) were intersected across all plates per condition.

### Data Normalization, Merging and Clustering

Batch effects due to technical noise were minimized using multiBatchNorm to downscale all batches (i.e. half-plates) to match the coverage of the least-sequenced batch. Feature selection was performed by selecting genes with the largest positive average biological component across all batches. PCA computation was performed on the selected set of genes across all batches using multiBatchPCA which ensures that each batch contributes equally to the definition of the PC space. Batches were then merged using fastMNN to detect mutual nearest neighbors (MNN) of cells in different batches to correct the values in each PCA subspace. Merging was performed using chosen variable genes selected among all expressed genes and detected within at least 5% of cells across all runs and conditions. Samples were first merged ‘by condition’ (all half-plates per condition) then ‘across conditions’ (E10.5 and E12.5). Seurat tSNE and UMAP values were calculated using fastMNN reduction with 1:50 dims. Clustering was obtained by random walks (cluster_walktrap, k=12) on a shared nearest-neighbor (SNN) graph (buildSNNGraph).

### Bioinformatic Analysis

Markers for each cluster were identified (findMarkers) using pairwise t-tests to keep genes that were differentially expressed in at least half of the pairwise comparisons with the other clusters (https://osca.bioconductor.org/marker-detection.html). Average expression per cluster was determined using AverageExpression on clusters that were subset using Seurat. Pseudo-time analysis was inferred using slingshot and the EMP cluster was used as a starting point. Genes associated with pseudo-time were obtained by adjusting the p-values using the Bonferroni method. Correlation to the Mouse Cell Atlas was made by comparison with a provided reference matrix and exclusion of cell types that matched less than 10 cells (https://github.com/ggjlab/scMCA). Normalized data from Nestorowa et al. were provided by the Hemberg lab (https://hemberg-lab.github.io/scRNA.seq.datasets, https://github.com/hemberg-lab/scRNA.seq.datasets/) and cells were integrated using fastMNN. Overlay of cells from each cluster onto two-dimensional flow cytometry plots was performed by defining a bi-exponential scale using ggplot2. Code will be made available on GitHub.

## ACKNOWLEDGEMENTS

We are indebted to Baptiste Saudemont and Yann Loe-Mie for critical advice, training, reagents and quality control for the MARS-Seq pipeline. We would like to thank Rebecca Gentek, Marc Bajénoff (CIML in Marseille) and Kémy Ade for providing and acquiring exploratory data for *Cdh5*^*CreERT2*^ experiments, Pascal Dardenne and Yvan Lallemand for handling and injections of mice, Sébastien Bastide and Caroline Kaiser for advice on scRNA-Seq analysis with Seurat, Tobias Weinberger and Rebeca Ponce Landete for isolation of adult tissues, Anne Dejean for transfer of the *Cdh5*^*CreERT2*^ strain and other members of the Gomez Perdiguero and Cumano groups for critical discussions and feedback. We appreciate the support and advice of the Cytometry and Biomarkers (UTECHS CB) platform (Sophie Novault and Sandrine Schmutz), the Center of Bioinformatics, Biostatistics and Integrative Biology (C3BI), the Institute Pasteur Single Cell Initiative (Heather Marlow) and the Animalerie Centrale. This work was funded by the Institut Pasteur, the CNRS, Revive (Investissement d’Avenir; ANR-10-LABX-73) and the European Research Council ERC investigator award (2016-StG-715320 to E.G.P.) E.G.P. also acknowledges financial support from the Fondation Schlumberger (FRM FSER 2017) and the Emergence(s) program from Ville de Paris (2016 DAE 190). L.I. was supported by a PhD fellowship from the Revive Labex and L.F. by a Florence Gould-Pasteur Foundation fellowship.

## Notes

### Competing Interest Statement

The authors have declared no competing interest.

