## Supplemental Figures for "Overlapping Definitive Progenitor Waves Divide and Conquer to Build a Layered Hematopoietic System"

### Supplemental Figure S1

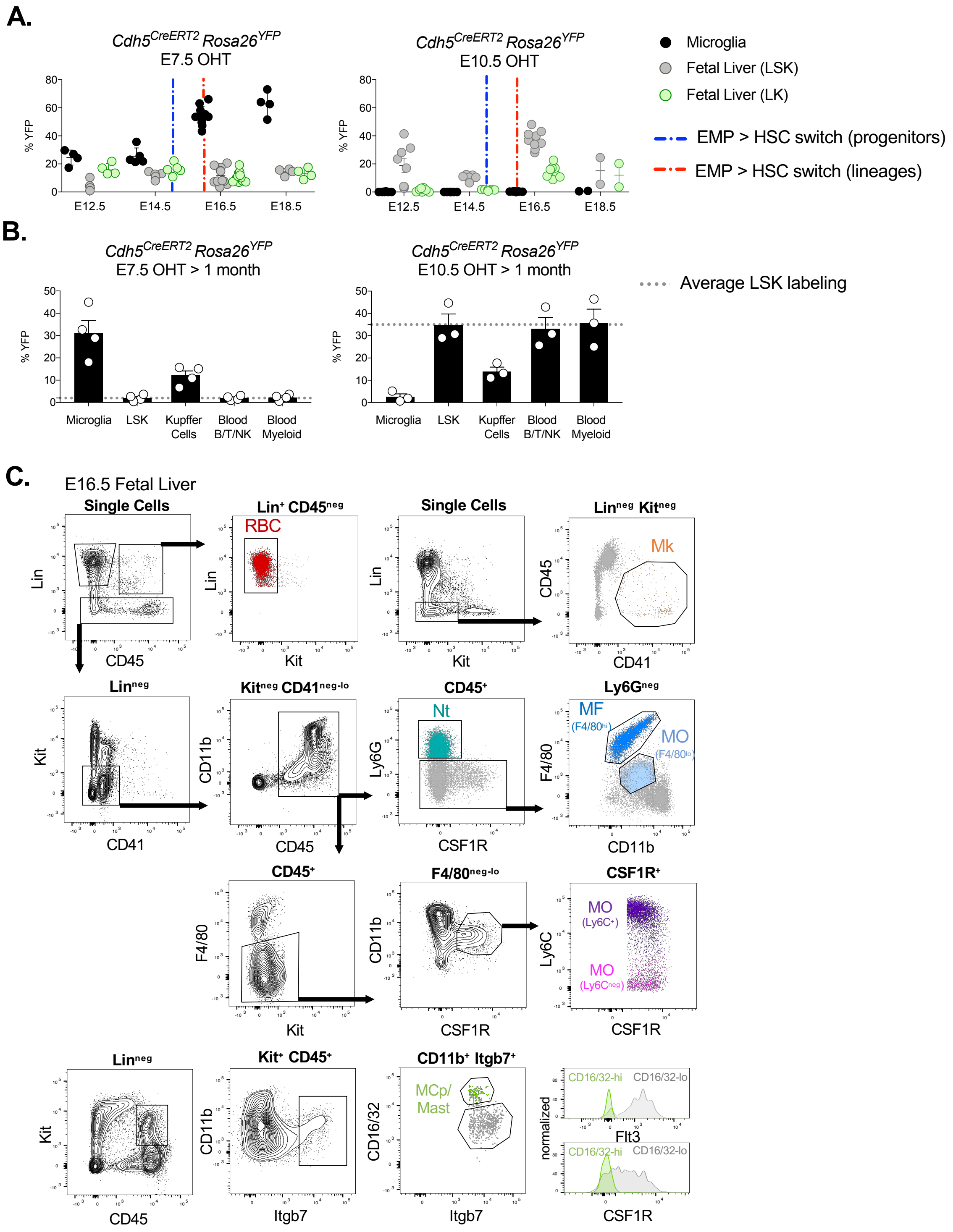

**Figure S1. EMP and HSC *in vivo* labeling using *Cdh5*$^{CreERT2}$ *Rosa26*$^{YFP}$.** **A.** Efficiency of *Cdh5*$^{CreERT2}$ *Rosa26*$^{YFP}$ pulse chase labeling among microglia (CD45+ F4/80+) with E7.5 OHT injection or among LSK (Lin$^{neg}$ Sca1$^{+}$ Kit$^{+}$) with E10.5 OHT injection. YFP labeling among LK (Lin$^{neg}$ Sca1$^{neg}$ Kit$^{+}$) progenitors is shown for blood and fetal liver. **B.** Percent YFP labeling among microglia, LSK, Kupffer cells (CD45+ F4/80$^{hi}$), circulating lymphoid (CD45$^{+}$ Lin$^{+}$ and CD19$^{+}$/CD3e$^{+}$/Nk1.1$^{+}$) and myeloid (CD45$^{+}$ Lin$^{neg}$ and Ly6G$^{+}$/Ly6C$^{+}$) cells is shown at one month of age following *in utero* labeling by OHT injection. Comparison among populations to the average LSK labeling (red dotted line) indicates partial replacement of EMP-derived Kupffer cells by HSC-derived cells after birth. **C.** Gating strategies for flow cytometry analysis of myeloid (Nt, neutrophil; F4/80$^{hi}$ MF, macrophage; F4/80$^{lo}$ MO, monocyte) and erythroid (RBC, red blood cell;, Mk, megakaryocyte) lineages. Data are represented as mean ± SEM.

#### Supplemental Figure S2

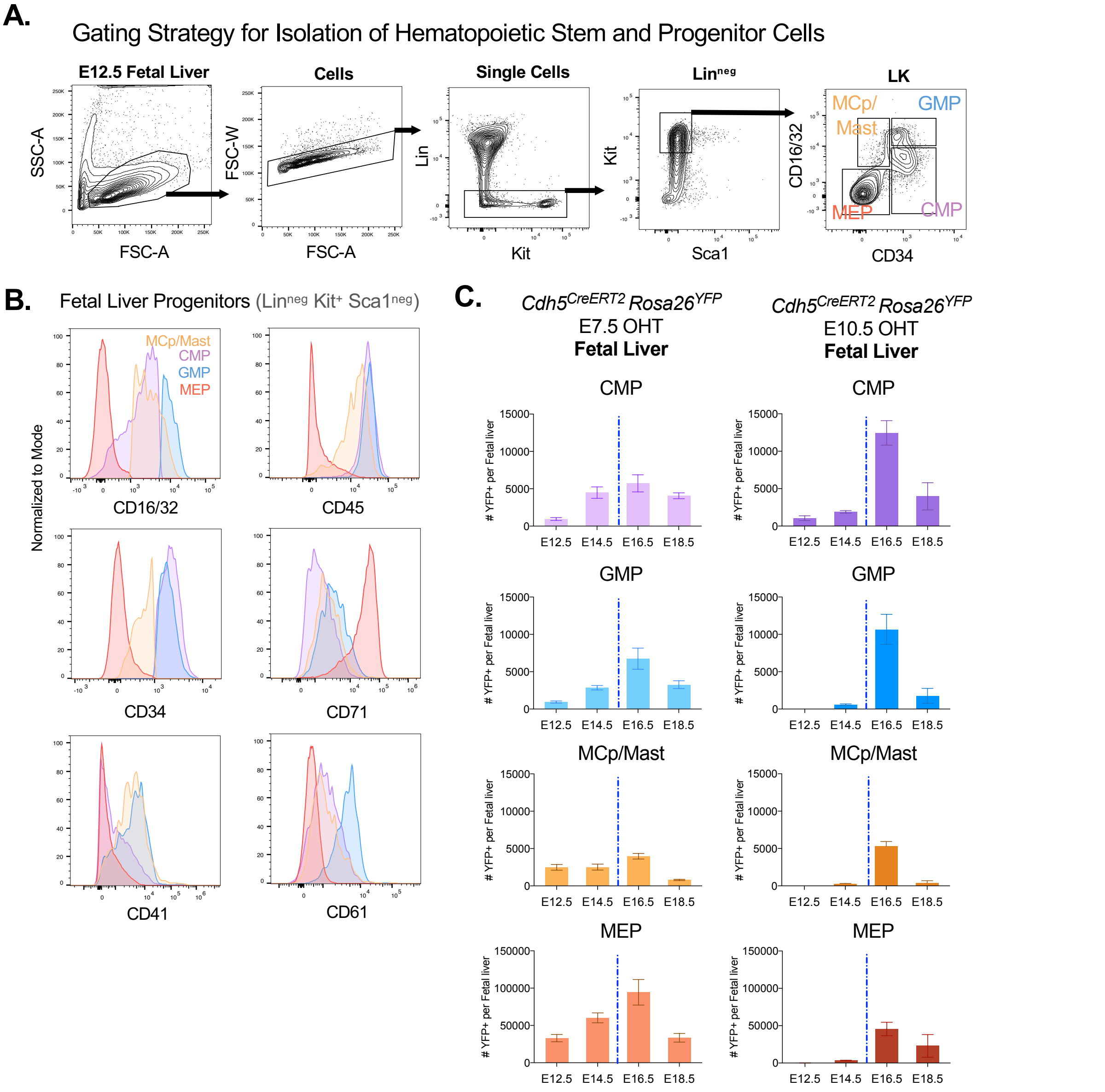


**Figure S2. Gating Strategy for Classification of Fetal Liver HSPC. A.** Gating strategy for classification of hematopoietic stem and progenitor cells (HSPC) into CMP (common myeloid progenitor), GMP (granulocyte-monocyte progenitor), MCp/Mast (mast cell progenitor) and MEP (megakaryocyte-erythrocyte progenitor) by flow cytometry using CD16/32 and CD34 cell surface markers. **B.** Example from E12.5 fetal liver validating the gating of CMP, GMP, MCp/Mast and MEP based on differential intensities of additional markers CD45, CD71, CD41 and CD61. **C.** Numbers of YFP$^{+}$ progenitors per fetal liver from *Cdh5$^{CreERT2}$ Rosa26$^{YFP}$* pulse chase labeing with E7.5 OHT (left) or E10.5 OHT (right). Data are represented as mean ± SEM.

#### Supplemental Figure S3

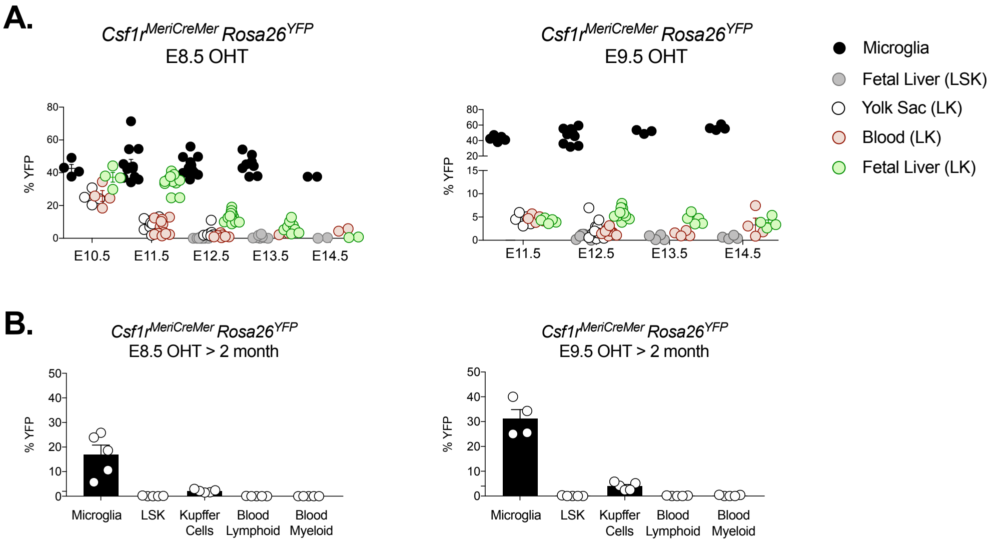


**Figure S3. Readout of *in vivo* EMP labeling using *Csf1r$^{MeriCreMer}$ Rosa26$^{YFP}$*. A.** Efficiency of *Csf1r$^{MeriCreMer}$ Rosa26$^{YFP}$* pulse chase labeling with E8.5 OHT or E9.5 OHT injection to label EMP was determined by percent of YFP$^+$ cells among microglia (CD45+ F4/80+). YFP labeling among LK (Lin$^{neg}$ Sca1$^{neg}$ Kit$^+$) progenitors is shown for yolk sac, blood and fetal liver. **B.** Percent YFP labeling among microglia, LSK, Kupffer cells (CD45+ F4/80$^{hi}$), circulating lymphoid (CD45$^+$ Lin$^+$ and CD19$^+$/CD3e$^+$/Nk1.1$^+$) and myeloid (CD45$^+$ Lin$^{neg}$ and Ly6G$^+$/Ly6C$^+$) cells is shown at two months of age following *in utero* labeling by E8.5 OHT or E9.5 OHT injections.

### Supplemental Figure S4

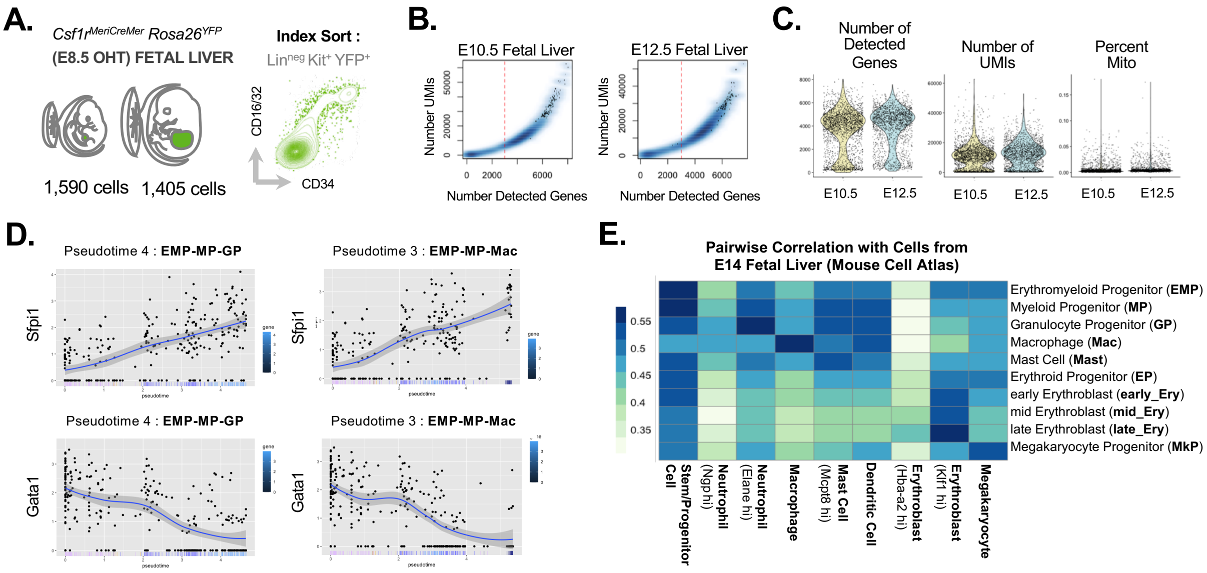


**Figure S4. scRNA-seq of EMP-derived Progenitors in the Fetal Liver.** **A.** Strategy for scRNA-seq of Lin$^{neg}$ Kit$^{+}$ YFP$^{+}$ fetal liver cells isolated from *Csf1r$^{MeriCreMer}$ Rosa26$^{YFP}$* embryos pulsed with E8.5 OHT using MARS-Seq technology. Cells were index sorted to capture cell surface intensity of CD34 and CD16/32 for retrospective correlation with single cell transcriptiomes. **B.** Cells were filtered by eliminating those with less than 3,000 detected genes that correlated with low UMI count. **C.** Cells were further filterd to remove those with greater than 5% mitochondrial fraction. **D.** *Sfpi1* and *Gata1* expression are shown along myeloid pseudotime trajectories. **E.** Heatmap of the correlation coefficients between average gene expression per cluster compared with E14 fetal liver cells annotated in the Mouse Cells atlas.

#### Supplemental Figure S5

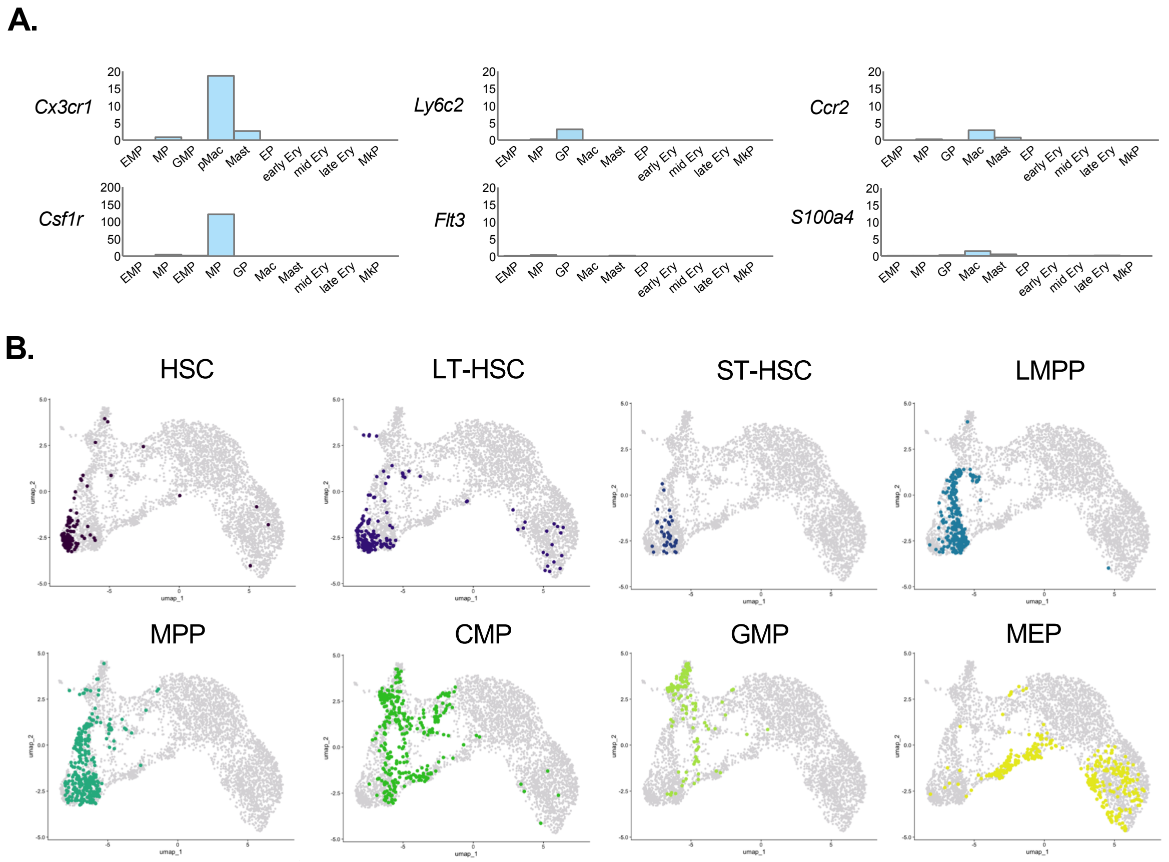


**Figure S5. Integration of EMP-derived Fetal Liver Progenitors with adult HSC/HSPC. A.** Average expression of genes among clusters. Selected genes are expressed in macrophages and pre-macrophages (*Cx3cr1, Csf1r*) or monocytes and monocyte progenitors (*Ly6c2, Flt3, Ccr2, S100a4*). **B.** UMAP representation of fetal liver EMP-derived progenitors integrated with previously published adult HSC/HSPC (Nestorowa et al., 2016). Clusters from the Nestorowa dataset are highlighted individually for visualization purposes.

### Supplemental Figure S6

A.

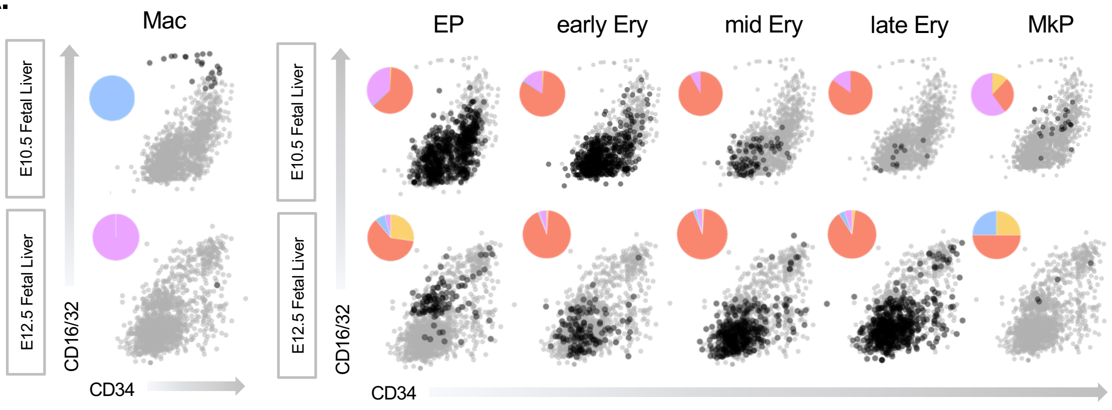


**Figure S6. Index Sorting of Mac and Erythroid Progenitors.** **A.** Index sorting classification of Mac and Ery cells from scRNASeq data using conventional CD16/32 versus CD34 gating to define CMP, GMP and MEP progenitor subsets for retrospective plotting of individually sequenced cells.
